# Effects of fibroblast growth factor 2 on muscle precursor cells from mouse limb and extraocular muscle

**DOI:** 10.1101/2025.08.17.670754

**Authors:** Austin J. Winker, Laura L. Johnson, Ria Jadhav, Catherine Nguyen, Elizabeth Hitch, Linda K. McLoon

## Abstract

Fibroblast growth factor 2 (FGF2) is known to play a role in skeletal muscle development and growth. We examined two populations of myogenic precursor cells for their responses to FGF2 *in vitro* using both extraocular and limb skeletal muscle. Fluorescence-activated cell sorting (FACS) was used to isolate two different populations of myogenic precursor cells, the EECD34 cells [positive for CD34, and negative for Sca1, CD31, and CD45] and PAX7-positive cells, from tibialis anterior and extraocular muscles of mice. These cells were cultured and treated with either proliferation or differentiation media in the absence or the presence of FGF2, followed by assays to determine its effects on proliferation and differentiation. These cells were also assessed for expression of fibroblast growth factor receptor (FGFR) 1, FGFR2, and FGFR4. Both the EECD34 cells and the PAX7-positive cells responded to FGF2 with significantly increased proliferation. Both myogenic precursor cell populations showed increased differentiation in the presence of FGF2, but also showed decreased rates of fusion into multinucleated myotubes in this *in vitro* system relative to control cells. FGF2 has pleiotropic effects on skeletal muscles. Contrary to the literature, FGF2 did not inhibit differentiation, but did appear to decrease fusion into multinucleated myofibers *in vitro*. These results provide a potential mechanism for reduction in myofiber number and size in the extraocular muscles in individuals with Apert syndrome, where FGF receptor 2 mutations maintain the receptor in an activated state.

## INTRODUCTION

Skeletal muscle contains resident precursor cells that enable repair and/or regeneration after injury or disease [1,2]. The first of these cells to be described were the satellite cells, which largely sit between the sarcolemma of the muscle fibers and the basal lamina surrounding each individual myofiber [3]. These cells are responsible for the ability of skeletal muscle to regenerate in muscle disease or after injury due to their ability to become activated, proliferate, and repair or regenerate new muscle fibers [4]. A number of studies have investigated the factors that control the proliferation and differentiation of muscle precursor cells in limb skeletal muscle. A wide range of neurotrophic and growth factors have been tested for their ability to stimulate proliferation and/or differentiation [5,6]. For example, fibroblast growth factor 2 (FGF2) promoted proliferation of satellite cells *in vitro* while insulin-like growth factor 1 (IGF1) promoted differentiation in satellite cells isolated from adult limb skeletal muscle from a number of animal species [5]. These factors also play a role in muscle regeneration and repair after injury or in the presence of skeletal muscle disease [7,8]. FGF2 has been shown to be particularly important in the maintenance of adult muscles, as its expression is up-regulated in muscle hypertrophy [9] and its absence in adult mice results in gait disturbances and muscle dysfunction [10].

Recent studies demonstrated that the myogenic precursor cells in skeletal muscles are not the same in different muscles [11]. Even within the well-described PAX7-positive satellite cells, a well-accepted marker for quiescent myogenic precursor cells found in all skeletal muscles [12], there is biochemical and functional heterogeneity [13,14]. In the extraocular muscles, there are differences in the populations of myogenic precursor cells when compared to limb skeletal muscles. In an earlier study we calculated this by both the total number of nuclei per total myofiber number in tissue sections and as the number of PAX7-positive cells per muscle weight quantified via flow cytometry [15]. We found that the density of PAX7-positive satellite cells in extraocular muscles is significantly elevated in number compared to limb skeletal muscles [15]. We performed several studies demonstrating that these PAX7-positive cells are preferentially spared after injuries compared to those in limb skeletal muscle, such as after irradiation [16] and are preferentially retained in the mdx mouse model of muscular dystrophy [17,18,19]. We identified a second large population of myogenic precursor cells, the EECD34 cells, identified as CD34-positive but negative for Sca1, CD31, and CD45 [18]. CD34 is commonly used as a marker to identify and purify muscle precursor cells, although common to many stem cells types [20]. Its expression is linked to the capacity for proliferation during skeletal muscle regeneration [21]. In the extraocular muscles, we showed using FACS analysis that this EECD34 population was 70% PITX2-positive [22]. In the absence of PITX2 expression in mice, the extraocular muscles lose many of their unique properties, including decreased expression of the extraocular muscle specific myosin heavy chain (MyHC) isoform (*myh13*) and loss of the *en grappe* endplates [23,24]. Based on isolation of these two populations using flow cytometry in our laboratory [22], only 3.65% of the Pitx2-positive cells expressed PAX7. This population is highly elevated in the extraocular muscles, and preferentially returns after 18Gy irradiation in both a mouse model of muscular dystrophy and in control mice, but not in limb muscle [16]. This suggests that besides being elevated in number, these cells from extraocular muscles have properties that may be different than this same population in limb skeletal muscle.

In addition to the differences in overall myogenic precursor cell numbers in normal extraocular muscles, extraocular muscles display a significant level of myonuclear addition into existing myofibers throughout life, as demonstrated using bromodeoxyuridine labeling methods [25,26]. These data were confirmed in other laboratories using a PAX7-reporter mouse, which showed extensive myonuclear addition in the normal mouse adult extraocular muscles compared to limb skeletal muscles [27]. Collectively, these data suggest that the myogenic precursor cells in the extraocular muscles are likely to respond differently to neurotrophic and growth factors than similar cells isolated from limb skeletal muscle.

We isolated EECD34 cells and PAX7-positive cells from mouse extraocular muscles and limb skeletal muscles, specifically the tibialis anterior, and examined their expression of fibroblast growth factor receptors 1, 2, and 4 (FGFR1, FGFR2, FGFR4) immunohistochemically *in vitro*. In separate cultures, isolated EECD34 cells and PAX7-positive cells from mouse extraocular muscles and limb skeletal muscles were examined for their rates of proliferation and differentiation *in vitro* after the addition of fibroblast growth factor 2 (FGF2) compared to control cultures to determine if there were differences in their behavior that might shed light on how the extraocular muscles are able to maintain their relatively high numbers of these myogenic progenitor cells. The response of the myogenic precursor cells to FGF2 might provide an explanation of their differential responses to short term compared to long term exogenously added FGF2 in rabbit extraocular muscles *in vivo* [28], where after short-term treatment the extraocular muscles showed increased muscle force generation, but with sustained treatment (3 months) muscle force generation was significantly decreased [28]. These results were interesting, as previous studies in limb skeletal muscle treated with FGF2 showed increased numbers of PAX7-positive cells, promoting regeneration [7,10,29,30]. Additionally, the results of these *in vitro* experiments may shed light on the phenotype of individuals with genetic mutations that involve FGFR expression, such as Apert syndrome [31], where we have shown significant reduction in overall muscle size in the extraocular muscles from individuals with this genetic disorder [32].

## METHODS

### Mice

Two mouse genotypes were used for cell isolation experiments. All studies were approved by the University of Minnesota Institutional Animal Care and Use Committee at the University of Minnesota. All studies adhered to the principles for use of animals in research set by the National Institutes of Health. For EECD34 cell isolation, we used C57BL/6 mice purchased from The Jackson Laboratory (Bar Harbor, ME). For PAX7-positive cell isolation, we used a PAX7-reporter mouse. *Pax7-Cre^+/-^; tdTomato-stop^fl/fl^* mice were generated by crossing *tdTomato^fl/-^* mice (JAX strain #: 007909) with *Pax7^cre/ERT2^* mice (JAX strain #: 017763). Resulting offspring were bred to produce *Pax7-Cre^+/-^; tdTomato-stop^fl/fl^* mice for experimental use, and the colony was maintained with *Pax7-Cre:^+/-^;tdTomato-stop^fl/fl^* (male) and *Pax7-Cre:^-/-^;tdTomato-stop fl/fl* (female) breeding pairs. All mice were housed by Research Animal Resources at the University of Minnesota Twin Cities. All mice used for these experiments were euthanized by tank CO_2_ asphyxiation at 3 months of age. Reporter expression was turned on in *Pax7-Cre^-/+^; tdTomato-stop fl/fl* mice via a once-daily/five-day injection series of tamoxifen. Tamoxifen was prepared at a concentration of 20 mg/ml in corn oil and injected intraperitoneally at a dose of 75 mg tamoxifen per kg body weight. A waiting period of two weeks after the fifth injection was required for full activation of tdTomato expression.

### Isolation of Myogenic Precursor Cells

#### PAX7-Positive Cells

Extraocular muscle (EOM) and tibialis anterior (TA) muscles were collected and pooled from 3 mice into chilled Dulbecco’s Modified Eagle Medium (DMEM) (Gibco, Billings, MT). EOM samples were placed into 1x phosphate buffered saline (PBS) containing collagenase B (Gibco) at 10 mg/ml, dispase II (Gibco) at 2.4 units/ml, and CaCl_2_ at 2.5 mM. Prior to placement in the collagenase/dispase solution, TA samples were chopped quickly with a razor blade to aid in digestion. Samples were incubated at 37⁰C, 5% CO_2_, for 90 minutes, and briefly removed every 25 minutes for pipette dissociation. The resulting cell suspension was filtered through a 70 µm filter and pelleted at 1400 rpm for 10 minutes at 4°C. After pouring off the supernatant, the pellet was resuspended in PBS-based sorter buffer containing 95% v/v PBS (Gibco, catalog # 10010023), 2.5% v/v fetal bovine serum (FBS) (Sigma Aldrich, St. Louis MO) and 2.5% v/v HEPES buffer (Sigma-Aldrich). The resuspended cells were filtered through a 40 µm filter into 5 ml FACS tubes and centrifuged at 1400rpm at 4°C for 5 minutes. Supernatant was discarded, and cells were resuspended in 2 ml of sorter buffer containing SYTOX Blue Dead Cell Stain (Invitrogen, catalog #S11348; Carlsbad, CA) at a 1:10,000 dilution. All work was performed in a sterile cell culture hood. Cells were kept on ice between steps.

#### EECD34 Cells

Samples used for EECD34 cell collection were prepared similarly to PAX7-positive cells with the following changes [22.33]. After filtering through a 70 µm filter, pelleting, and resuspending, the cells were incubated in the dark on ice for 45 minutes with fluorescent antibodies targeting CD34 (5 µl/100 µl sorter buffer, BV421 rat anti-mouse CD34, catalog #562608, BD Biosciences, Franklin Lakes, NJ), Sca1 (2 µl/100 µl sorter buffer, FITC rat anti-mouse Ly-6A/E, catalog #553335, BD Biosciences,), CD45 (2 µl/100 µl sorter buffer, PE/Cyanine7 rat anti-mouse CD45, catalog #103114, BioLegend, San Diego, CA), and CD31 (5 µl/100 µl sorter buffer, APC rat anti-mouse CD31, catalog #551262, BD Biosciences). Cells were filtered through at 40 µm filter, centrifuged, and resuspended in sorter buffer containing SYTOX Orange Nucleic Acid Stain (catalog #S11368, Invitrogen). All work was done in a sterile cell culture hood. Cells were kept on ice between steps.

#### FACS

PAX7-positive and EECD34 cells were collected from the EOM and TA samples with a BD FACSAria II at the University of Minnesota Flow Cytometry Resource Center. For PAX7-positive cell sorting, tdTomato^+^/SYTOX Blue^-^ cells were collected. For EECD34 cell sorting, CD34^+^/Sca1^-^/CD45^-^/CD31^-^/SYTOX Orange^-^ cells were collected. All cells were collected into 2 ml of proliferation media containing 95% DMEM, 10% fetal bovine serum, 10% horse serum, 0.5% chick embryo extract, and 1% penicillin/streptomycin.

#### Cell Culture and FGF2 Treatment

After cells were sorted, they were pelleted and resuspended in proliferation media to a concentration of 32 cells/µl. Thus, for all *in vitro* experiments, 250 µl of these cells were plated on Matrigel Membrane Matrix (Corning, catalog #CB-40234A, Corning, NY) coated Lab-Tek 8 Well Chamber Slides (catalog #62407-335, Corning) and kept in an incubator at 37⁰ C and 5% CO_2_. Every 48 hours, proliferation media was aspirated and replaced with fresh proliferation media. For the proliferation assay, at ∼30% confluency, cells were treated with recombinant mouse FGF2 (Bio-Techne, catalog #319-FB-025, Minneapolis, MN) at 10 ng/ml or 40 ng/ml in proliferation media. For the differentiation assay, when cells were at 30% confluency, roughly 4 days for EECD34 cells and 3 days for PAX7-positive cells, they were placed in low serum differentiation media supplemented with 10 ng/ml or 40 ng/ml of recombinant mouse FGF2 (Bio-Techne). The low serum media was aspirated and replaced with fresh media containing consistent amounts of FGF2 every 48 hours. The low serum media consisted of 94% v/v DMEM (Gibco), 5% v/v horse serum (R&D Systems, catalog #S12150, Minneapolis, MN), and 1% v/v penicillin-streptomycin (Gibco, catalog #CX30321). Control wells for both the proliferation and differentiation assays were supplemented with Dulbecco’s Phosphate-Buffered Saline (PBS) only (Gibco, catalog #CX30294).

#### Immunofluorescent Staining of FGF Receptor on PAX7 and PITX2 Expressing Cells

For both EECD34 cells and PAX7 cells isolated by FACS, an FGFR stain was performed at the same time point as the proliferation assay with FGF2 treatment. To start, the media was aspirated, and cells were fixed in 4% paraformaldehyde at room temperature for 15 minutes before being washed with 3% BSA w/v in PBS. They were permeabilized with 0.5% Triton X-100 (Sigma-Aldrich) in PBS for 10 minutes at room temperature. Cells were washed with 3% BSA w/v in PBS before being incubated with antibody to fibroblast growth factor receptor 1 (FGFR1), FGFR2, or FGFR4, as follows: anti-FGFR1 (1:500, #9740; Cell Signaling, Danvers, MA, USA), anti-FGFR2 (1:150; #23328; Cell Signaling), or anti-FGFR4 (1:100, #8562, Cell Signaling). For the secondary antibody, the cells were incubated in goat-anti-rabbit IgG PLUS AF488 (1:1000, A32731, Invitrogen) for 30 minutes. After a PBS rinse, they cells were counterstained with the nuclear marker Hoechst 33342 and coverslipped using Vectashield Vibrance anti-fade mounting medium (H-1700, Vector Labs., Newark, CA).

#### Cell Proliferation Assay

Still in their culture plates, the cells were incubated in proliferation media with 10 µM EdU for 1 hour at 37⁰C and 5% CO_2_ 24 hours following FGF2 treatment. Then, cells were fixed in 4% paraformaldehyde at room temperature for 15 minutes before being washed with 3% BSA w/v in PBS, and then permeabilized with 0.5% Triton X-100 (Sigma-Aldrich, catalog #T9284) in PBS for 10 minutes at room temperature. Cells were washed with 3% BSA w/v in PBS before being incubated with ClickiT EdU Cell Proliferation Kit reagents (Invitrogen, catalog #C10337) at room temperature for 30 minutes in the dark to label proliferating cells. After washing with 3% BSA w/v in PBS, cell nuclei were incubated with Hoechst 33342 (part of ClickiT kit) in PBS at 1:2000 dilution for 15 minutes at room temperature in the dark. Cells were washed in 3% BSA w/v in PBS before being cover-slipped with H1000 Vectashield Mounting Media (Vector Laboratories, catalog #NV9265087, Burlingame, CA). Cells were imaged on a Leica DM4000B Fluorescent Microscope (Leica, Wetzlar, Germany). A minimum of 10 images at 20x magnification were quantified per treatment well. Proliferating cell percentages from these images were averaged to determine the percent of cells in each category from the same biological replicate. These averages were used to calculate the percent of cells proliferating in a minimum of 4 independent replicates per treatment. Proliferating cell percentages were calculated for each image using the Bioquant Imaging System (Bioquant, Nashville, TN).

#### Cell Differentiation Assay

Once EOM cells reached ∼80% confluency, cells in low-serum media treated for FGF2 were assayed for their differentiation status. This usually occurred 1-2 days after the change to differentiation media with FGF2 for PAX7+ cells and 2 days after for EECD34 cells. Because TA cells generally grew slower than EOM and were not necessarily treated with FGF2 at the same time, TA cells were assayed on later days than EOM but on the same timeline as described here. For the assay, the media was aspirated, and wells were washed with PBS before being fixed in 4% paraformaldehyde in PBS. After washing in PBS, cultures were blocked in 10% horse serum for 30 minutes before use of avidin and biotin blocking solutions (Vector Laboratories, catalog #SP2001; Newark, CA) according to package instructions. Cells were incubated in desmin antibody (Dako, M0760, Troy, MI) at a 1:500 concentration for 1 hour at room temperature to label differentiating cells. The Vectastain Elite ABC-HRP kit (Vector Labs.) was used according to package instructions to enable colorimetric desmin staining when cells were consequently treated with diaminobenzidine (DAB). Slide chambers were removed, and slides were submerged in hematoxylin to visualize nuclei. The slides were coverslipped using H1000 Vectashield Mounting Media (Vector Labs.). Levels of cell differentiation were defined as follows: desmin-negative and thus non-differentiating with a single nucleus; desmin-positive and thus differentiating with a single nucleus; and desmin-positive differentiating with multiple nuclei. These populations were quantified using the Bioquant Imaging System (Bioquant, Nashville TN). Approximately 20 images at 20x magnification were quantified per treatment well. These were averaged to determine the percent of cells in each category from the same biological replicate. These averages were used to calculate the percent of cells in each differentiation state in a minimum of 4 independent replicates per treatment.

##### Statistical Analysis

All data are reported as means +/- SD. All statistical analyses were performed in Prism 9 (GraphPad, La Jolla, CA). One-way ANOVAs were performed to determine significance. If the data were significantly different, post-hoc Tukey’s multiple comparison tests were performed for TA and EOM data to determine significance between treatment groups. Statistical significance was defined as P<0.05.

## RESULTS

### FGF receptor (FGFR) expression

Upon reaching 30% confluence, both EECD34 cells and PAX7 FACS-isolated cells from wild type mice from extraocular muscle and TA were treated with FGF2 and 24 hours later were immunostained for their expression of the three FGFRs known to be expressed in skeletal muscle: FGFR1, FGFR2, and FGFR4 [6]. In the cultures of mononucleated EECD34 cells derived from either EOM or TA, 98.9% and 99.5% of these expressed FGFR1, respectively (Figure 1A,C and Figure 2A,C). The mononucleated PAX7-positive cells isolated from EOM were 98.3% positive for FGFR1 expression (Figure 2B). The PAX7-positive cells isolated TA were 100% positive for FGFR1 expression (Figure 2D).

**Fig 1.**
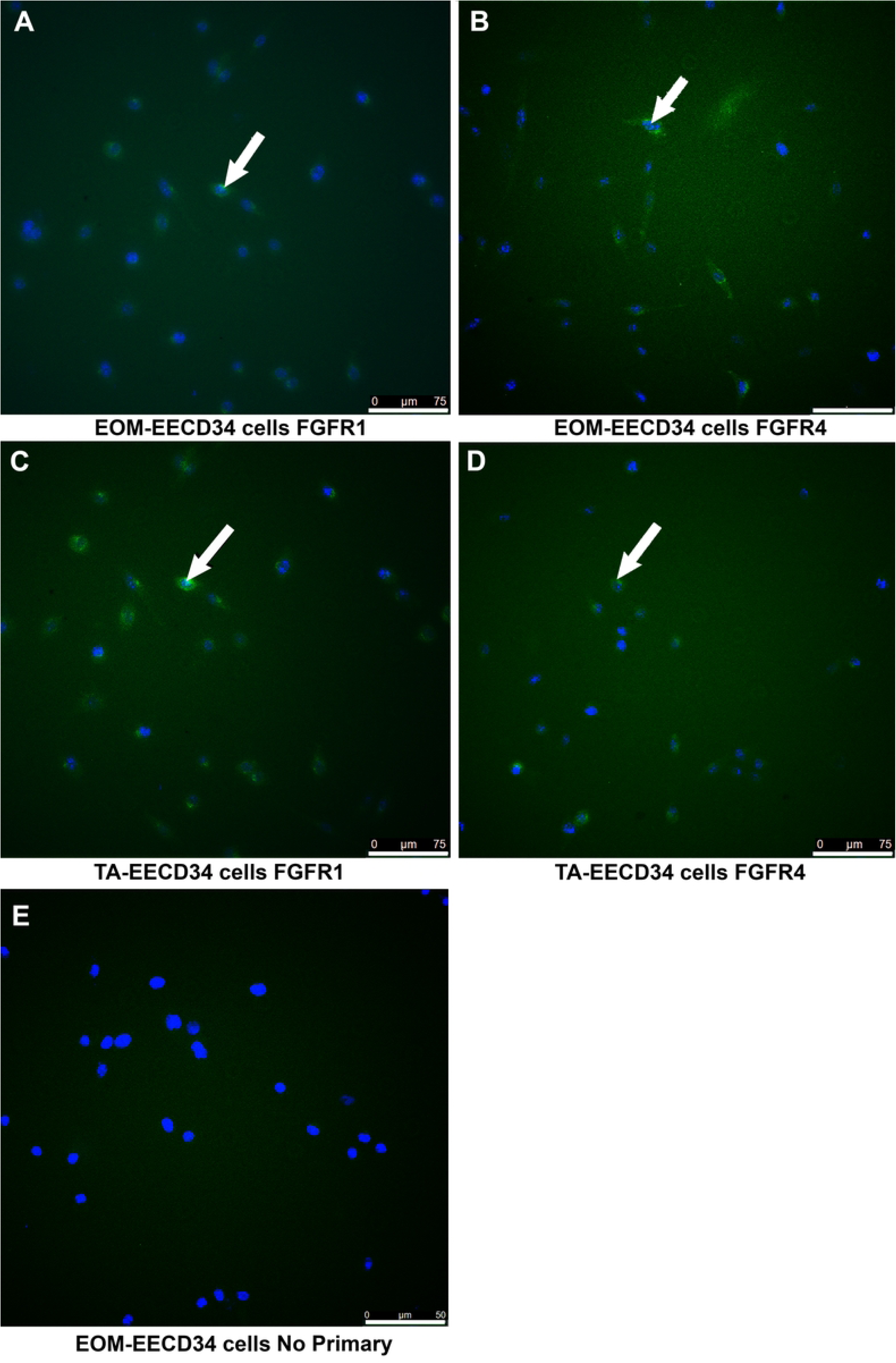
EECD34 cell isolated from EOM or TA were cultured and 24 hours after reaching 30% confluence and treatment with FGF2, assessed for their expression of fibroblast growth factor receptors (FGFR). (A) EECD34 cells isolated from EOM immunostained for FGFR1. (B) EECD34 cells isolated from EOM immunostained for FGFR4. (C) EECD34 cells isolated from TA immunostained for FGFR1. (D) EECD34 cells isolated from TA immunostained for FGFR4. (E) EECD34 cells isolated from EOM immunostained in the absence of primary antibody as a control for non-specific binding. Essentially all the cells were positive for FGFR1, and the majority of cells were positive for FGFR4. White arrows indicate FGFR-positive cells. A-D: Magnification bar is 75 µm. E: Magnification bar is 50 µm.

**Fig 2.**
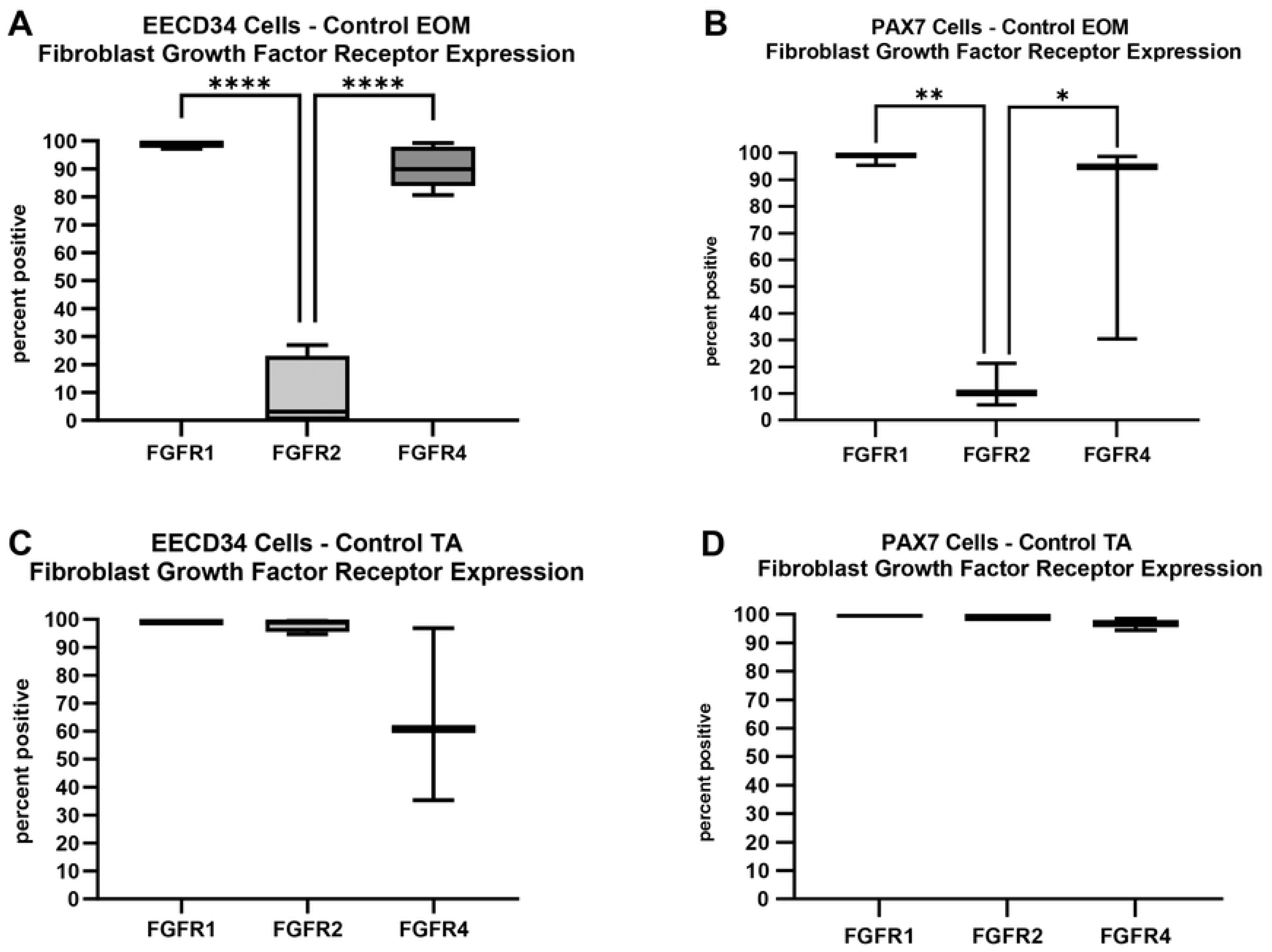
Quantification of FGFR expression in EECD34 cells and PAX7 cells immunostained after 3 days *in vitro*. (a) EECD34 cell isolated from EOM immunostained for FGFR1, FGFR2, and FGFR4. **** indicates p<0.0001. (B) PAX7-positive cells isolated from EOM immunostained for FGFR1, FGFR2, and FGFR4. ** indicates p<0.001. * indicates p<0.01. (C) EECD34 cells isolated from TA immunostained for FGFR1, FGFR2, and FGFR4. (D) PAX7-positive cells isolated from EOM immunostained for FGFR1, FGFR2, and FGFR4.

The EECD34 cells derived from EOM were only 9.9% positive for the expression of FGFR2 (Figure 2A). This expression level was significantly lower than for expression of FGFR1 (p=0.0001) and significantly lower than for the expression of FGFR4 (p=0.0001). In contrast, the cells derived from TA were 98.1% positive for the expression of FGFR2 (Figure 2C). Only 12.5% of the PAX7-positive cells isolated from EOM expressed FGFR2; this was significantly lower than expression of FGFR1 (p=0.009) or FGFR4 (p=0.04). In contrast, 99% of the PAX7-positive cells isolated from TA expressed FGFR2 (Figure 2B, D).

The expression of FGFR4 mirrored that of FGFR1. In the EECD34 cells derived from EOM, 90.7% expressed FGFR4 (Figure 2A); however, the EECD34 cells derived from TA showed a more variable expression, with 64.3% of the cells expressing FGFR4 (Figure 2C). In the cultures of PAX7-positve cells derived from EOM, 74.7% expressed FGFR4 (Figure 2B), while 96.6% of those derived from TA expressed FGFR4 (Figure 2D). Thus, the receptors responsible for FGF2 signaling were present on our cultured cells, with clear co-expression of several of the FGFR subtypes.

### EECD34 Cells - Proliferation

We examined the effect of FGF2 treatment of EECD34 cells on proliferation rate using EdU labeling *in vitro*. ANOVA analysis showed that FGF2 significantly increased the percent of proliferating cells (p<0.01). Post-hoc analyses showed increases from the control percentage of 49% to 61.5% for the 10ng/ml treatment (p=0.03) and to 62% for the 40ng/ml treatment (p=0.029) (Figure 3A-C,G). These represent percent increases of 25.4% and 26.5% respectively. For the treated EECD34 cells derived from TA, the ANOVA showed significance (p=0.003). Post-hoc t-tests showed that the 10 ng/ml treatment increased the percent proliferating cells from the control level of 24.8% to 41.43%, a significant increase of 67.4% (p=0.007). The 40 ng/ml treatment similarly increased the percent of proliferating cells to 39.3%, a significant increase of 58.6% (p=0.005) (Figure 3H).

**Fig 3.**
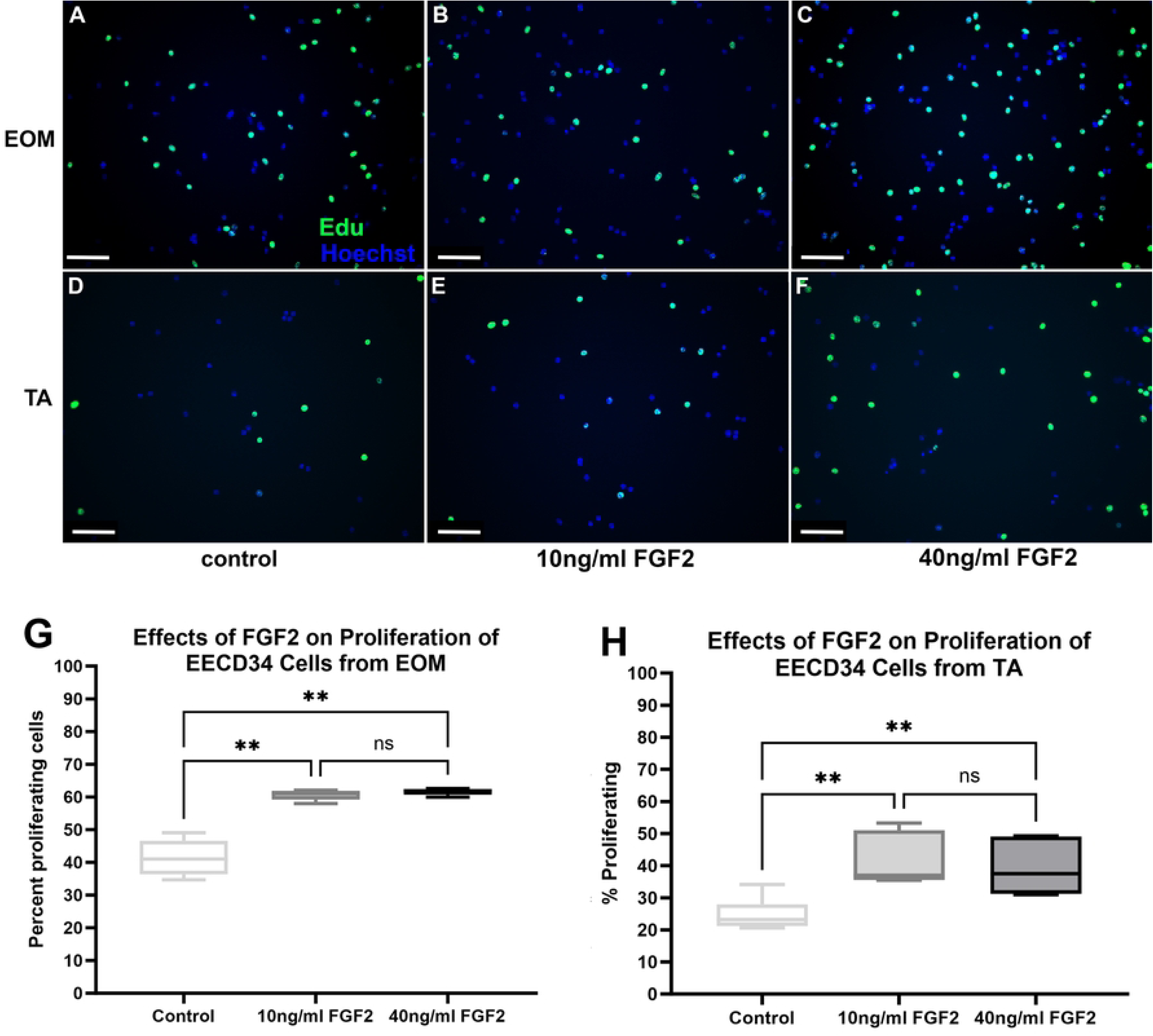
FACS-isolated EECD34 cells from extraocular muscles (EOM) (A-C) and tibialis anterior (TA) (D-F) immunostained for EdU (green) and Hoechst dye (blue). Grown for 3 days in high serum (20%) media to promote proliferation. (A,D) Untreated control cells, (B,E) cells treated with 10 ng/ml FGF2, and (C,F) cells treated with 40 ng/ml FGF2. Magnification bar is 75 µm. Quantification of the rate of proliferation of FACS-isolated EECD34 cells derived from EOM or TA treated with FGF2. (G) Quantification of EECD34 cells derived from EOM after 24 hours in FGF2-supplemented proliferation media. FACS-isolated EECD34 cells were cultured until approximately 30% confluent and then treated with either 10 ng/ml or 40 ng/ml FGF2 in proliferation media for 24 hours. Proliferation was determined by examining EdU uptake for 1 hour immediately following the 24-hour treatment. Immunohistochemistry was used to visualize the EdU-positive cells (green) compared to non-proliferating cells (blue). These data represent 8 independent replicates for each treatment. (H) Quantification of EECD34 cells derived from TA after 24 hours in proliferation media. FACS-isolated EECD34 cells were cultured, treated, and assayed in the same manner as the EOM EECD34 cells. Immunohistochemistry was used to visualize the EdU-positive cells (green) compared to non-proliferating cells (blue). These data represent eight independent replicates for each treatment. * indicates significant difference from control.

### EECD34 Cells - Differentiation

EECD34 cells from EOM were grown in differentiation media and treated with either 10 ng/ml or 40 ng/ml FGF2, and ANOVA analysis showed significant differences (p=0.0001) (Figure 4A-G). Post-hoc multiple comparison tests showed a significant decrease of desmin-negative cells when treated with 40 ng/ml FGF2 compared to controls (70.28% decrease, p=0.036), and a trend toward decreasing of desmin-negative cells when treated with 10 ng/ml FGF2 that was not significant (69.28% decrease; p=0.056). There were significant increases in the number of desmin-positive cells with single nuclei after both 10 ng/ml and 40 ng/ml treatment (23.84% increase; p=0.014; 25.5% increase, p=0.008), respectively. Interestingly, there was a significant decrease in rate of fusion (fusion index) in these differentiating cultures in the presence of 10 ng/ml or 40 ng/ml FGF2, decreasing by 62.9% (p=0.024) and 68.3% (p=0.02), respectively.

**Fig 4:**
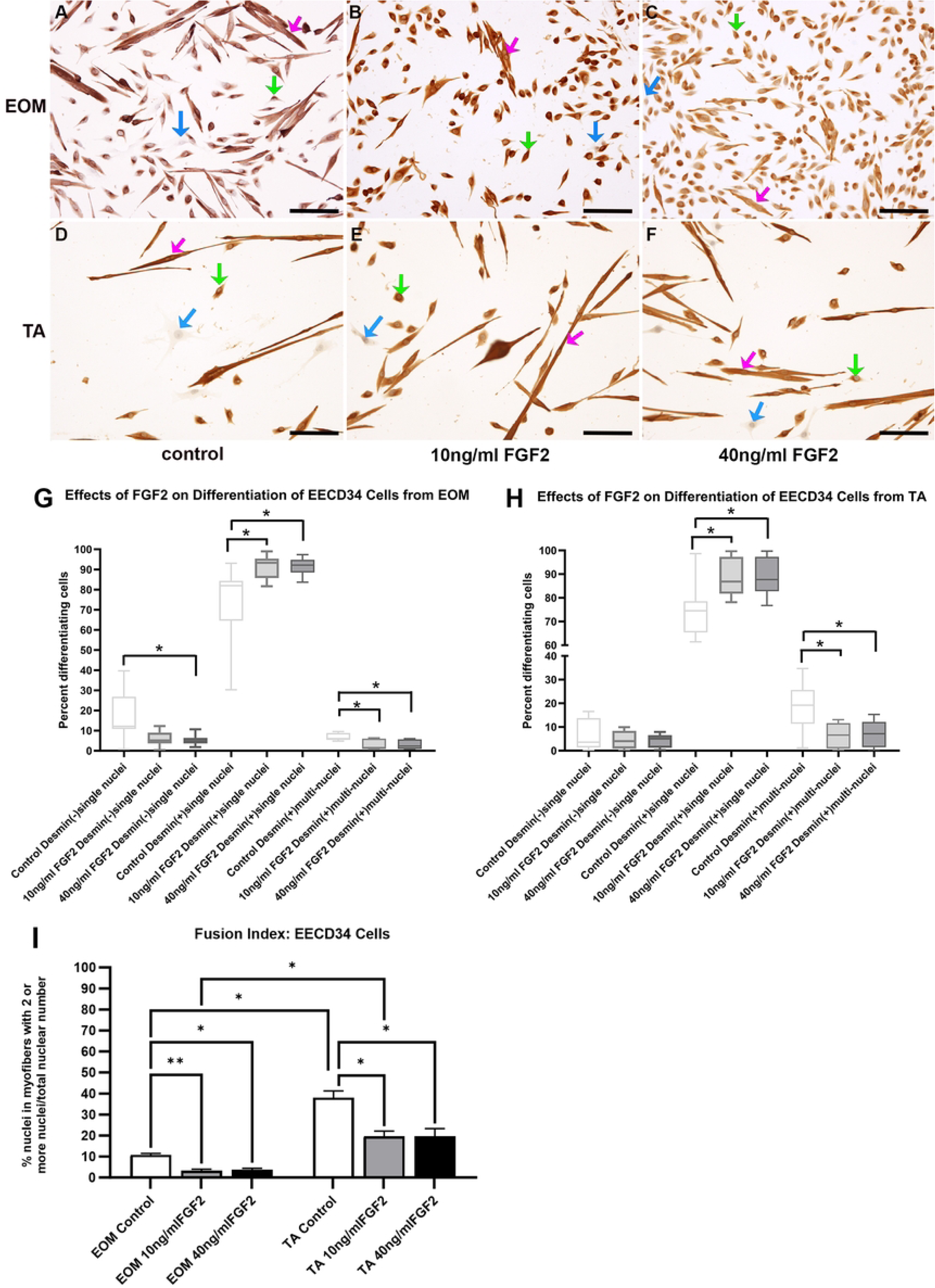
FACS-isolated EECD34 cells from EOM (A-C) and TA (D-F) immunostained for desmin (brown) and counterstained with hematoxylin (blue) grown in the low-serum medium to promote differentiation. (A,D) Untreated control cells, (B,E) cells treated with 10 ng/ml FGF2, and (C,F) cells treated with 40 ng/ml FGF2. Bar is 100µm. Blue arrow: undifferentiated cells; Green arrow: desmin-positive singly nucleated cells; Pink arrow: desmin-positive multiply nucleated cells. Quantification of FACS-isolated EECD34 cells derived from the EOM or TA after being placed in differentiation media, treated with FGF2 at 30% confluency and stained two days later. (G) FACS-isolated EECD34 cells from EOM were cultured and treated with either 10ng/ml or 40ng/ml FGF2 and compared to cells grown in differentiation media only. Two days later, the cells were fixed, and differentiation was determined by examining expression of desmin. Cells were categorized as desmin-negative, desmin-positive containing only one nucleus, or desmin-positive containing more than one nucleus. These data represent seven independent replicates. (H) FACS-isolated EECD34 cells from TA were cultured, treated, and assayed in the same manner as the EOM EECD34 cells. These data represent seven independent replicates. * indicates significant difference from control within the same treatment group. (I) Analysis of fusion index calculated as the number of nuclei with EECD34 cells that contain two or more nuclei as a percent of all nuclei in all cells counted. * indicates significance is at least p<0.04. ** indicates significance is at least p<0.009.

EECD34 cells from TA were grown in differentiation media and treated with either 10 ng/ml or 40 ng/ml FGF2; ANOVA analysis indicated significant differences (p=0.0001) (Figure 4H). Post-hoc multiple comparison tests showed that compared to the untreated controls, there were no significant differences in the number of desmin-negative cells after either of the FGF2 treatments (p=0.67; p=0.58, respectively). There were significant increases in the number of desmin-positive cells with single nuclei after both 10 ng/ml and 40 ng/ml treatment (18.9% increase, p=0.018; 18.7% increase, p=0.024, respectively). There was a significant decrease in rate of fusion in these differentiating cultures in the presence of 10 ng/ml or 40 ng/ml FGF2 compared to controls, decreasing by 65.4% (p=0.012) and 62.2% (p=0.02), respectively.

### PAX7-Positive Cells - Proliferation

FGF2 treatment of PAX7-positive cells derived from EOM was examined for its effect on proliferation rate (Figure 5). The ANOVA analysis demonstrated significant differences (p=0.002). Post-hoc analysis demonstrated that, in contrast to the EECD34 cells, both 10 ng/ml and 40 ng/ml of FGF2 resulted in a significant increase in the proliferating cell population. The 10 ng/ml treatment increased the percent proliferating cells from 30.7% to 52.2%, an increase of 70.1% (p=0.002). The 40 ng/ml treatment similarly increased the percent proliferating cells to 47.7% compared to the controls, an increase of 55.4% (p=0.012). The ANOVA analysis demonstrated that, in contrast to the EECD34 cells, FGF2 treatment of PAX7-positive cells isolated from TA were significantly different from control proliferation rates (p=0.002). Post-hoc analysis demonstrated that the 10 ng/ml treatment increased the percent proliferating cells from 34.2% to 47.3%, an increase of 38.3% (p=0.028). Despite a 25.1% increase in cell number after treatment with 40 ng/ml FGF2, this was not statistically significant (p=0.15).

**Fig 5.**
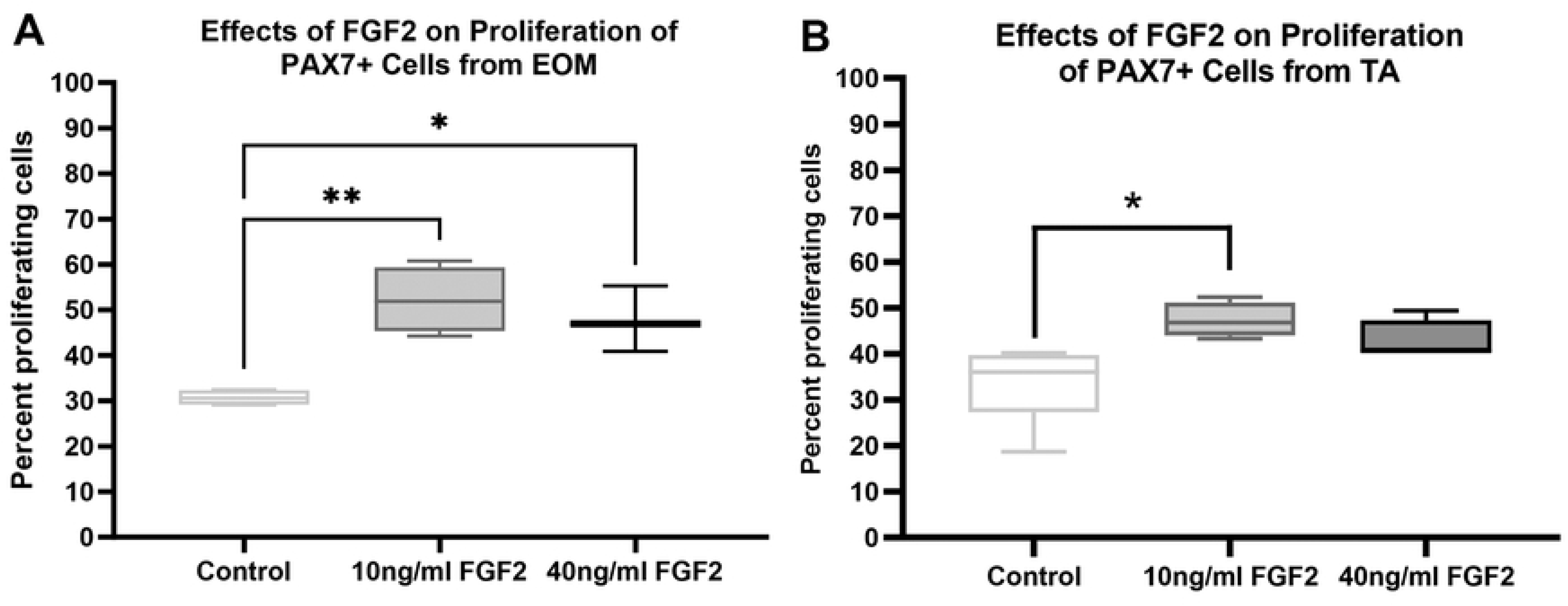
Quantification of the rate of proliferation of FACS-isolated PAX7-positive cells derived from EOM or TA treated with FGF2. (A) Quantification of PAX7-positive cells derived from EOM after 24 hours in proliferation media. FACS-isolated PAX7-positive cells were cultured until approximately 30% confluent and then treated with either 10 ng/ml or 40 ng/ml FGF2 in proliferation media for 24 hours. Proliferation was determined by examining EdU uptake for 1 hour immediately following the 24-hour treatment. Immunohistochemistry was used to visualize the EdU-positive nuclei (green) compared to all nuclei (blue). These data represent 4 independent replicates for each treatment. (B) Quantification of PAX7-positive cells derived from TA after 24 hours in proliferation media. FACS-isolated PAX7-positive cells were cultured until approximately 30% confluent and then treated with either 10 ng/ml or 40 ng/ml FGF2. Proliferation was determined by examining EdU uptake after 1 hour. Immunohistochemistry was used to visualize the EdU-positive cells (green) compared to all cells (blue). These data represent 4 independent replicates for each treatment. * indicates significant difference from control within the same treatment group.

### PAX7-Positive Cells - Differentiation

PAX7-positive cells from EOM were grown in differentiation media and treated with either 10 ng/ml or 40 ng/ml FGF2 (Figure 6A-G). ANOVA analysis showed a significant difference (p=0.0001). Post-hoc multiple comparison tests showed that there was a significant increase in the number of desmin-positive cells with single nuclei at both 10 ng/ml and 40 ng/ml doses, increasing from 92.18% to 97.7% (p=0.0005) and 98.14% (p=0.0003), respectively. Similarly, there was a reduction in fusion compared to the control cells, with a decrease from the control at 7.71% down to 2.3% in cells treated with 10 ng/ml FGF2 (p=0.0005) and 1.8% in cells treated with 40 ng/ml FGF2 (p=0.0003). PAX7-positive cells derived from TA showed similar responses to those isolated from the EOM. ANOVA analysis showed a significant difference (p=0.0001) (Figure 6H). Post-hoc multiple comparison tests showed that there was a significant increase in the number of desmin-positive cells with single nuclei at both 10 ng/ml and 40 ng/ml relative to the controls. The percent of these cells went from 82.3% in controls cells up to 92.94% in cells treated with 10 ng/ml FGF2 (p=0.02) and 94.69% in cells treated with 40 ng/ml FGF2 (p=0.014). Similarly, there was a reduction in fusion compared to the control cells, with a decrease from the control at 17.15% down to 5.9% in cells treated with 10 ng/ml FGF2 (p=0.019) and 5.3% in cells treated with 40 ng/ml FGF2 (p=0.014).

**Fig 6.**
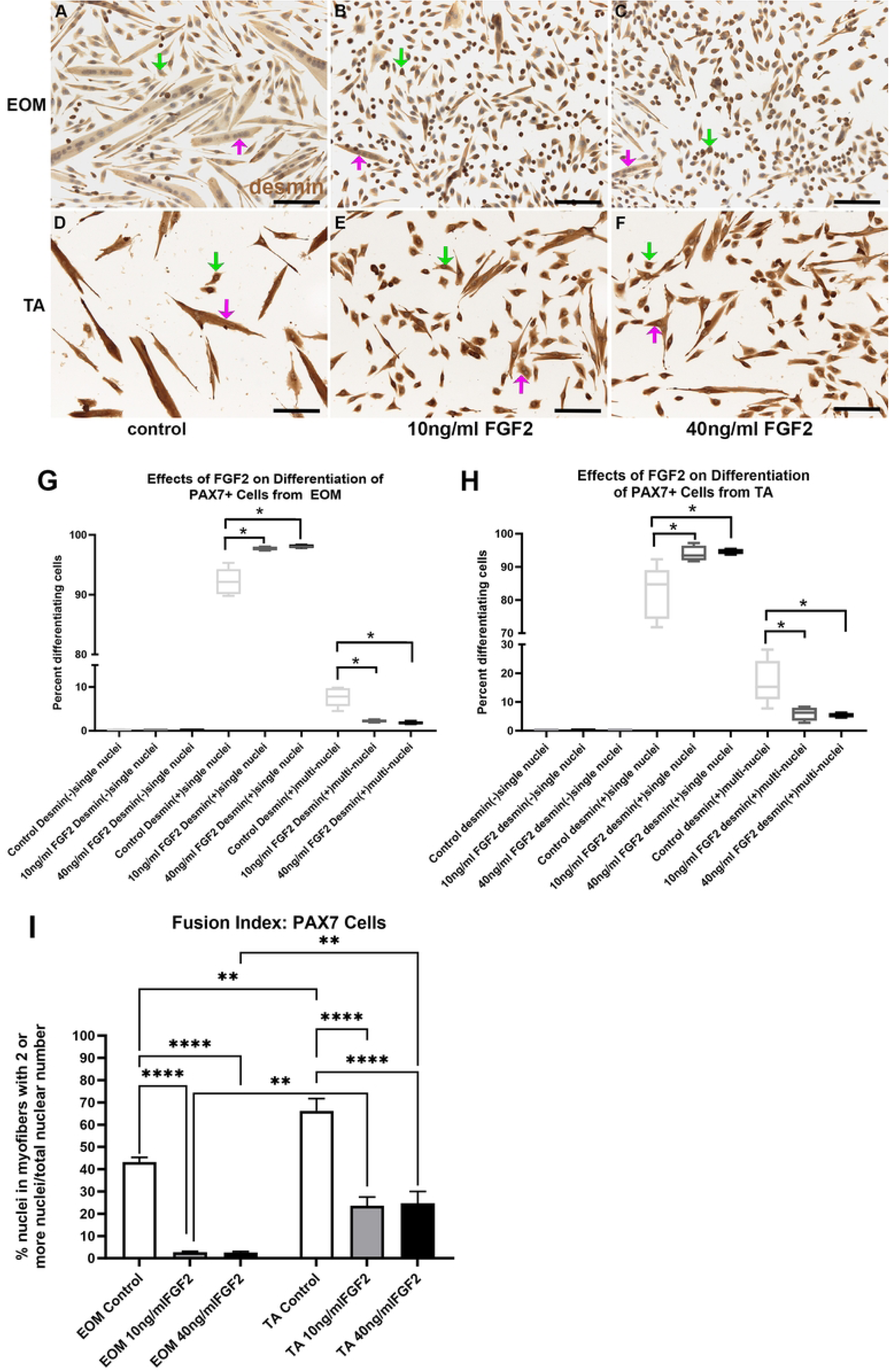
PAX7-positive cells isolated from EOM (A-C) and TA (D-F) immunostained for desmin (brown) and counterstained with hematoxylin (blue) grown in the low-serum medium to promote differentiation. (A,D) Untreated control cells, (B,E) cells treated with 10 ng/ml FGF2, and (C,F) cells treated with 40 ng/ml FGF2. Green arrow: desmin-positive singly nucleated cells; Pink arrow: desmin-positive multiply nucleated cells. Bar is 100 µm. Quantification of FACS-isolated PAX7-positive cells derived from the EOM (G) or TA (H) after being placed in differentiation media and treated with FGF2 at 30% confluency, and stained two days later. (G) FACS-isolated PAX7-positive cells from EOM were cultured and treated with either 10 ng/ml or 40 ng/ml FGF2 and compared to cells grown in differentiation media only. Two days later, the cells were fixed, and differentiation was determined by examining expression of desmin. Cells were categorized as desmin-negative, desmin-positive containing only one nucleus, or desmin-positive containing more than one nucleus. These data represent 4 independent replicates. (H) FACS-isolated Pax7-positive cells from TA were cultured, treated, and assayed in the same manner as the EOM Pax7-positive cells. Cells were categorized as desmin-negative, desmin-positive containing only one nucleus, or desmin-positive containing more than one nucleus. These data represent 5 independent replicates. * indicates significant difference from control within the same treatment group. (I) Analysis of fusion index calculated as the number of nuclei with Pax7 cells that contain two or more nuclei as a percent of all nuclei in all cells counted. ** indicates significance is at least p<0.008. **** indicates significance is at least p<0.0001.

A fusion index for the PAX7 cells was calculated based on number of nuclei within desmin-positive cells that contained two or more nuclei compared to the total nuclei in all cells (Figure 6I). This analysis demonstrated that in the cultures of PAX7 cells derived from EOM there were significant decreases of the number of nuclei within multinucleated fibers in FGF2 treated cultures compared to control cultures, with decreases in 10ng/ml FGF2 cultures of 93.7% [p=0.0001] and 94.2% in cultures treated with 40 ng/ml FGF2 (p=0.0001). Decreased numbers of nuclei within multinucleated fibers was also see in cultures of PAX7 cells derived from TA, with decreases in cultures treated with 10 ng/ml FGF2 by 64.5% [p=0.0001] and decreases in cultures treated with 40 ng/ml FGF2 by 62.8% (p=0.0001). As seen with the EECD34 cells, the percent of nuclei within multinucleated fibers was significantly greater in control cultures of PAX7 cells derived from TA than EOM (34.5% increase, p=0.0034). In the presence of 10 ng/ml FGF2, the EOM-derived cultures showed significantly lower numbers of nuclei within multinucleated cells compared to the TA derived cultures (88.5% decrease, p=0.0008). Similarly, there was a decreased number of nuclei within multinucleated cells in the cultures treated with 40 ng/ml FGF2 derived from EOM compared to those from TA, with an 89.8% decrease (p=0.005), supporting the hypothesis that there are intrinsic differences between what are ostensibly the same cells derived from EOM compared with those from TA.

## DISCUSSION

Our data demonstrate that *in vitro,* FGFR1, FGFR2, and FGFR4 are expressed on both the EECD34 cells and PAX7-positive populations of myogenic precursor cells derived from adult mouse extraocular muscle, similar to patterns found using Northern blot analysis in developing human limb skeletal muscle [34]. Thus, the receptors for producing responses by these cells to exogenously added FGF2 were in place.

The EECD34 cells are approximately 70% PITX2-positive in EOM and about 65% PITX2-positive from leg muscle [22] and represent a population of myogenic precursor cells particularly elevated in the extraocular muscles. Based on our flow cytometric cell isolations of PAX7 and PITX2-immunostained cells derived from normal genetically-unaltered mice followed by cytospin isolation, only 3.65% of PIXT2-positive cells labeled directly *ex vivo* co-expressed PAX7 [22]. Thus, in our hands, the EECD34 cells and the PAX7 cells represent different populations. The EECD34 cells isolated from adult mouse EOM muscle showed an increased rate of proliferation in the presence of FGF2 *in vitro*, similar to studies showing that FGF2 promotes the proliferation of isolated satellite cells *in vitro* [35,36]. It is also known that skeletal muscle satellite cells can express both FGF1 and FGF2 [37]. This supports experimental evidence that despite being isolated under the same parameters, defined by their expression of this set of specific precursor cell markers — CD34^+^/Sca1^-^/CD31^-^/CD45^-^ — they have some similar properties as the “same” cells derived from the TA, despite the comparatively unique properties of the EOM.

Similarly, the PAX7-positive cells isolated from EOM and TA showed a significantly increased rate of proliferation in the presence of added FGF2 *in vitro*. This agrees with a large number of previous studies, where isolated satellite cells – which are largely PAX7-positive – responded to FGF2 *in vitro* with increased rates of proliferation [5,38]. Direct intramuscular injection of FGF2 also was shown to result in increased proliferation within the treated limb skeletal muscles, with short-term increases in Ki67-positive and PAX7-positive cells [39]. It is interesting to note a complexity in the response of extraocular muscle to FGF2. One week after injection of FGF2 directly into an extraocular muscle in adult rabbits there was a significant increase in the number of PAX7-positive cells within the treated muscle [28]. However, after sustained delivery for 3 months of FGF2 in adult rabbit extraocular muscle, there was a significant decrease in satellite cell numbers and significantly reduced myofiber cross-sectional areas [28]. In FGF2 knockout mice, significant gait disturbance and decreases in muscle size were present [10], supporting the hypothesis that FGF2 is critical in maintaining the pool of muscle precursor cells needed for maintenance of normal muscle function over time.

A number of recent studies have examined the myogenic precursor cell populations in the extraocular muscle with conflicting results. Studies have demonstrated, in agreement with our bromodeoxyuridine pulse-labeling studies [25,26,40–42], that there is a continual, albeit slow, process of replication of these cell populations in the extraocular muscles [27] and that these cells appear to have greater and more long-lasting proliferative potential throughout life [19,21,43,44]. These properties appear to be part due to the environment of the niche [45,46] and part due to properties intrinsic to the extraocular muscles [19,46,47]. In contrast to these studies, a recent study using a number of transgenic mice described the PITX2-positive and PAX7-positive cell populations to be non-proliferative in the adult, in contrast to previous studies showing proliferation and myofiber fusion in many skeletal muscles [27,48] but agree that the extraocular muscles contain a higher content of PAX7-positive cells than limb skeletal muscles [49]. One potential explanation for this discrepancy could relate to transgenic mouse modifications in development that might alter the potential fates of the myogenic precursor cell populations in the adult mouse. Future studies will need to address this difference.

Using desmin as a marker for differentiation, FGF2 treatment of EECD34 and PAX7-positive cells *in vitro* resulted in a significant increase in the percent of differentiating cells, as identified by desmin expression, as well as a likely concomitant decrease in the number of desmin-negative undifferentiated cells. These results contrast with many previous studies showing that FGF2 significantly decreased differentiation *in vitro* and *in vivo* [5,7,36,50]. However, these sorts of pleiotropic effects are commonly seen with a variety of neurotrophic and growth factors. For example, insulin-like growth factor-1 can promote both the proliferation and differentiation of muscle precursor cells [51,52]. FGF2 also has been shown to have these pleiotropic effects [6,28].

Interestingly, despite the increase in the percentage of desmin-positive cells, for both the EECD34 cells and the PAX7-positive cells isolated from extraocular muscles and leg muscle, there was a decrease in the desmin-positive multinucleated cells. This suggests that while differentiation significantly increased, there was a concomitant reduction in the ability of these differentiating cells to fuse. An earlier study showed that high levels of FGF2 were found in myogenin-positive cells, which indicate that they were differentiating but also singly-nucleated [53]. This lack of fusion is interesting in light of several studies suggesting this also occurs when FGF2 levels are increased within skeletal muscles. Using FGF2 embedded in microspheres, injection into injured leg muscle resulted in increased myogenic precursor cell proliferation but no change to overall muscle force generation [54]. The ability of FGF2 to increase functional muscle mass was tested using FGF2 collagen scaffold implants after volumetric muscle loss, and while it resulted in increased myoblast proliferation, it was unsuccessful in increasing myofiber density [55]. In another study, short-term treatment of skeletal muscle with FGF2 resulted in some improvement in number of regenerated myofibers in the mdx mouse muscular dystrophy model, but the effect was largely due to enhanced replication of satellite cells in the treated muscles [7,29]. Also, local injection of FGF2 into a previously denervated thyroarytenoid muscle within the larynx again resulted in increased numbers of proliferating cells as well as increased numbers of MyoD-positive cells within the treated muscles, suggesting that FGF2 had the potential to decrease atrophy in the denervated muscles [39]. Similar results were obtained in aging laryngeal muscles, with some evidence that differentiation was promoted by these treatments [39,56]. Long-term functional changes after these treatments were not assessed in these two studies. We previously showed significantly different effects of short-term and long-term treatment with FGF2 [28]. While direct injection of FGF2 into a superior rectus muscle of an adult rabbit resulted in increased numbers of PAX7-positive cells and increased muscle force, one-month and three-month sustained treatment with FGF2 resulted in a significant decrease in the numbers of PAX7-positive cells and a significant decrease in force generated by the treated superior rectus muscles. These results are interesting in that they provide a potential mechanism for the small extraocular muscles sizes seen in individuals with Apert syndrome, which is also supported by the present study [32]. In Apert syndrome, in the vast majority of cases there are one of two mutations in the gene for FGFR2 [57,58]. These gain-of-function mutations result in increased affinity for the FGF ligand, maintaining prolonged action even when FGF levels are limited [59]. Similarly, other craniosynostosis disorders that result in strabismus, such as Crouzon syndrome and Pfeiffer syndrome, have gain-of-function mutations in FGFR2 [60]. The effect on extraocular muscles and the resultant strabismus is not surprising, as human fetal extraocular muscles express FGFR2 [61]. Thus, this *in vitro* study serves as a potential mechanism for the effects of prolonged signaling of FGFR2.

The apparent inhibition of fusion in the FGF2 treated cells was surprising, as FGF2 has not been linked to the control of myoblast fusion. This result supports our observation that after sustained treatment with FGF2, the extraocular muscles generated less force, and the myofibers were smaller than normal – presumably by further decreasing fusion of muscle precursor cells into existing fibers [28]. Interestingly, FGF2 is involved in controlling myoblast migration during development [62] and *in vitro* [63,64], which in turn could result in decreased fusion. Future studies will need to determine if this is the case. Interestingly, transforming growth factor-beta (TGFβ) also results in decreased myoblast fusion in limb skeletal muscle [65,66]. Expression levels of TGFβ appear to be quite low in normal extraocular muscles [67,68], but is expressed at elevated levels in strabismic muscles [68,69]. When we treated adult rabbit EOM with TGFβ, we found that the treated rabbit superior rectus muscles had significantly decreased mean myofiber cross-sectional areas from the already small fibers found in the extraocular muscles, and these muscles generated less force – similar to what was seen after extended FGF2 treatment [28,70]. These studies suggest that modulation of many different neurotrophic factors can result in similar outcomes when attempting to modify extraocular muscle structure and function.

With skeletal muscles, there are a large number of neurotrophic and growth factors whose expression overlap in normal adult, developing, and diseased or injured muscles. In a controlled tissue engineering experiment, after accelerating satellite cell activation and proliferation with FGF2, sequential addition of insulin-like growth factor 1 (IGF1) was a potent inducer of differentiation [71]. In the extraocular muscles, sequential expression of FGF2 and IGF-1 was observed in a time series shortly after an experimental recession in rabbit model [69], again suggesting that multiple growth factors play a complex, interdependent role in maintaining normal function in the extraocular muscles and in modulating the responses within the muscles after strabismus treatment. Our previous studies showed that sustained treatment of rabbit superior rectus muscles with FGF2 resulted in muscles that were smaller and produced less force than controls. In individuals with overacting muscles causing strabismus, sustained FGF2 treatment has the potential to improve alignment by weakening the appropriate muscle.

### Conclusions

In summary, in tandem with satellite cells isolated from leg skeletal muscle, the EECD34 myogenic precursor cells derived from EOM, although representing a potentially more multipotent population [18], respond to FGF2 *in vitro* with increased proliferation. Contrary to many previous studies, both the EECD334 and PAX7 cell populations responded with increased levels of differentiation, as identified with desmin expression, in the presence of added FGF2. However, cell fusion was reduced significantly, suggesting that FGF2 influences the ability of these cells either to move or to recognize each other. Further studies are needed to understand this observation. These studies provide a potential mechanistic explanation for the observed decreases in muscle size and number seen in the extraocular muscles in individuals with Apert syndrome [32], which is characterized by prolonged binding of FGF and activation of the FGFR2 [59].

## Acknowledgements

Supported by NIH EY15313 (LKM), NRT-UtB: Graduate Training Program in Sensory Science (AJW), T32EY025187 (AJW), and the Minnesota Lions Gift of Sight.

This manuscript will be a chapter in the PhD thesis of Austin Winker.

